# Combining images and anatomical knowledge to improve automated vein segmentation in MRI

**DOI:** 10.1101/152389

**Authors:** Phillip G. D. Ward, Nicholas J. Ferris, Parnesh Raniga, David L. Dowe, Amanda C. L. Ng, David G. Barnes, Gary F. Egan

## Abstract

**Purpose:** To improve the accuracy of automated vein segmentation by combining susceptibility-weighted images (SWI), quantitative susceptibility maps (QSM), and a vein atlas to produce a resultant image called a composite vein image (CV image).

**Method:** An atlas was constructed in common space from 1072 manually traced 2D-slices. The composite vein image was derived for each subject as a weighted sum of three inputs; a SWI image, a QSM image and the vein atlas. The weights for each input and each anatomical location, called template priors, were derived by assessing the accuracy of each input over an independent data set. The accuracy of venograms derived automatically from each of the CV image, SWI, and QSM image sets was assessed by comparison with manual tracings. Three different automated vein segmentation techniques were used, and ten performance metrics evaluated.

**Results:** Vein segmentations using the CV image were comprehensively better than those derived from SWI or QSM images (mean Cohen’s d = 1.1). Sixty permutations of performance metric and automated segmentation technique were evaluated. Vein identification improvements that were both large and significant (Cohen’s d>0.80, p<0.05) were found in 77% of the permutations, compared to no improvement in 5%.

**Conclusion:** The accuracy of automated venograms derived from the composite vein image was overwhelmingly superior to venograms derived from SWI or QSM alone.

## 1. Introduction

Mapping cerebral veins using magnetic resonance (MR) images has until recently been technically challenging. Cerebral venograms are increasingly important for advancing our knowledge of cerebral vascularisation, oxygenation, metabolism and studies of cerebrovascular topology. The use of venograms in clinical research applications is growing rapidly, including for quantifying oxygen saturation (Fan et al., 2014), measuring the metabolic rate of oxygen consumption (Rodgers et al., 2016) and analyzing possible fMRI confounders (Vigneau-Roy et al., 2014).

Traditional vein imaging techniques require invasive contrast agents, have potential arterial confounds, and are limited to the large vessels, due to the reduced volume and flow of smaller cerebrovasculature segments. However, magnetic susceptibility provides an intrinsic contrast mechanism that is exquisitely sensitive to the presence of iron, particularly deoxygenated iron-rich haemoglobin proteins within red blood cells, making susceptibility techniques very useful for imaging small as well as large veins. The magnetic susceptibility of blood is modulated by oxygen (Pauling and Coryell, 1936), which facilitates the separation of arteries and veins, whilst providing a mechanism to quantify oxygen saturation (Fan et al., 2014).

Susceptibility-weighted imaging (SWI) and quantitative susceptibility mapping (QSM) are MR techniques based on magnetic susceptibility that provide a non-invasive method of imaging the cerebral veins. QSM and SWI derive contrast from gradient-recalled echo (GRE) phase information and have been applied to stroke, multiple sclerosis, cerebrovascular disease, and examined in clinical and preclinical studies (Fan et al., 2015; Fujima et al., 2011; Goodwin et al., 2015; Jain et al., 2010; Li et al., 2013; Liu and Li, 2016; Rodgers et al., 2013; Santhosh et al., 2009). The way in which SWI and QSM process the phase information is very different.

SWI multiplies a non-linear mapping of high-pass filtered GRE phase with the GRE magnitude image, compounding the effects of signal cancellation from incoherent signals within each voxel and phase accumulation due to local sources of magnetic susceptibility (Haacke et al., 2004). Non-local sources are also included, such as the extravascular phase information, resulting in the magnification of small veins. The presentation of non-local sources, and non-linear mapping, generates a non-quantitative image best suited to radiological interpretation.

QSM estimates the magnetic susceptibility distribution directly by inverting the magnetic field information captured in the phase image (Marques and Bowtell, 2005; Salomir et al., 2003). Mathematically, QSM involves a linear system inversion that is ill-posed and requires regularization or fitting (Li et al., 2015; Liu et al., 2016; Wang and Liu, 2015; Wharton et al., 2010). QSM has the benefits of being quantitative and is designed to resolve extravascular field effects, leaving only local sources of magnetic susceptibility contrast.

The differing approaches (QSM and SWI) have unique image contrasts, and each have their own vein-like confounders. SWI images, for instance, do not distinguish between signal cancellation due to venous blood, and low concentrations of free protons (Haacke et al., 2004). The lack of distinction is problematic when analyzing veins which reside near non-vein low signal structures, such as in the vicinity of the tentorium and in the interhemispheric fissure (due to the falx cerebri). Both SWI and QSM also suffer different artefacts, such as cruciform artefacts in QSM images.

As neither QSM nor SWI isolate blood signal intrinsically, unlike spin-labelling or contrast agent-based techniques, venous voxels within the brain must be identified before the veins can be analysed. The process of identifying venous voxels in the brain, or vein segmentation, produces a vein mask (or venogram) that can then be used to extract the vein signal from an image, or examined directly for topographic analysis.

A number of algorithms for automatic segmentation of blood vessels in the body have been proposed, including shape-driven, intensity-driven and hybrid approaches (Lesage et al., 2009). A common approach in the analysis of SWI and QSM data is to employ a preliminary filtering step, such as Hessian-based filtering (Frangi et al., 1998), before applying a simple threshold classification method (Vigneau-Roy et al., 2014). Recent work has combined Hessian-based filtering into a segmentation framework with diffusion techniques to overcome noise and low vein visibility (Bazin et al., 2016; Manniesing et al., 2006). Statistical modeling of spatial relations has also been proposed to improve continuity and smoothness in vein segmentation (Bériault et al., 2014; Ward et al., 2017b).

The previously mentioned work focused upon SWI or QSM, and did not attempt to extract information from both images. Methods have been proposed that merge SWI with QSM (Ward et al., 2015), and R2* maps (Monti et al., 2015). Both approaches were globally homogeneous, i.e., they combined two images without consideration for anatomical location. As SWI and QSM have differing image contrasts, and artefacts that are specific to anatomy, it is possible a superior segmentation could be achieved if the method for combining the two images was sensitive to location.

Prior anatomical knowledge has recently been incorporated into a vein segmentation technique to reduce false positives in specific brain regions (Bériault et al., 2015). However, this approach was limited to specific deep-brain regions, it did not directly address boundaries between tissue types and neural structures, and it was hand-tuned.

There are two anatomical factors that contribute to vein segmentation accuracy. The first is vein anatomy, i.e., expected vein occurrence, size and shape at an anatomical location. The second is image contrast, i.e., expected tissue signal relative to vein signal, which is specific to SWI and QSM. In this study, these two factors are exploited to improve cerebrovenous contrast and subsequent vein segmentation accuracy. We propose a vein identification and segmentation method that is based on a locally varying combination of SWI and QSM contrast which is informed by known vein anatomy in specific neuroanatomical structures. The proposed method derives a single composite vein image (CV image) that incorporates the strengths of SWI and QSM, with the anatomical knowledge of a vein atlas.

The CV image is generated from three input images (SWI, QSM and atlas) that are combined using a weighted-sum. The weights are derived from template priors that capture the location-specific venous contrast of the three input images throughout the brain. Separate vein atlases and template priors were calculated for each subject within the study from an independent sample of the cohort to ensure data independence. Future applications of the technique would use a single template prior and atlas calculated from the entire cohort. The CV image was compared to SWI and QSM images for the purpose of vein segmentation using automated techniques. Segmented CV images were compared with segmented SWI and QSM images using a broad array of accuracy measures and three automated segmentation techniques.

## 2. Methods

All procedures were reviewed and approved by the local ethics committee. Informed consent was obtained from all volunteers. The code and data used in this study have been available to the public using github and figshare respectively (Ward et al., 2016a, 2017).

### 2.1 Data Acquisition

Ten healthy volunteers were scanned using a 3T Siemens Skyra with a 32-channel head and neck coil (6 females, mean age 56.2 years, standard deviation 25.2). The protocol was a single echo, flow-compensated, gradient-recalled echo (GRE) sequence (TE=20ms, TR=30ms, flip angle=15^o^, voxel=0.9×0.9×1.8mm anisotropic, matrix 256×232×72). Four of the subjects were acquired with a smaller voxel size (voxel=0.9mm isotropic, matrix=256×232×160). A T1-Weighted MPRAGE scan was also acquired and used for registration purposes (TE=2ms, TR=2300ms, TI=900ms, voxel=1.0mm isotropic, matrix=240x256x192, flip angle=9^o^). All registration was performed using FNIRT and FLIRT from the FSL toolkit (https://fsl.fmrib.ox.ac.uk/)(Jenkinson et al., 2012).

For all subjects, raw k-space data for the GRE acquisition was saved for each coil and retrospectively reconstructed to generate phase and magnitude images. Individual coil phase images were processed to remove phase wraps and background phase shifts using Laplacian unwrapping (Li et al., 2014) and V-SHARP (Wu et al., 2012). QSM maps were computed using LSQR in the STI-Suite (Li et al., 2014). The SWI images were taken directly from the scanner console.

Six of the ten subjects were healthy elderly subjects recruited for the ASPirin in Reducing Events in the Elderly (ASPREE) clinical trial (Group, 2013) and scanned at baseline as part of the ASPREE NEURO sub-study (S. A. Ward et al., 2016).

### 2.2 Manual vein tracing

A mask containing venous voxels was created for each subject by manually labelling voxels as vein or non-vein, using FSLView (Jenkinson et al., 2012). Tracing was performed by author PW under the supervision of author NF (a clinical radiologist). The venous voxels were initially identified based upon SWI contrast and prior anatomical knowledge, in transverse reconstructions of the 3D SWI acquisition, and refined by reference to sagittal and finally coronal reformats. Initial SWI-only masks were then overlaid side-by-side on SWI and QSM images for editing. Editing was performed slice by slice in the sagittal plane, followed by the axial and finally the coronal plane. An example ground-truth venogram is shown in Figure **1**. In total 1072 transverse slices were examined.

**Figure 1.**
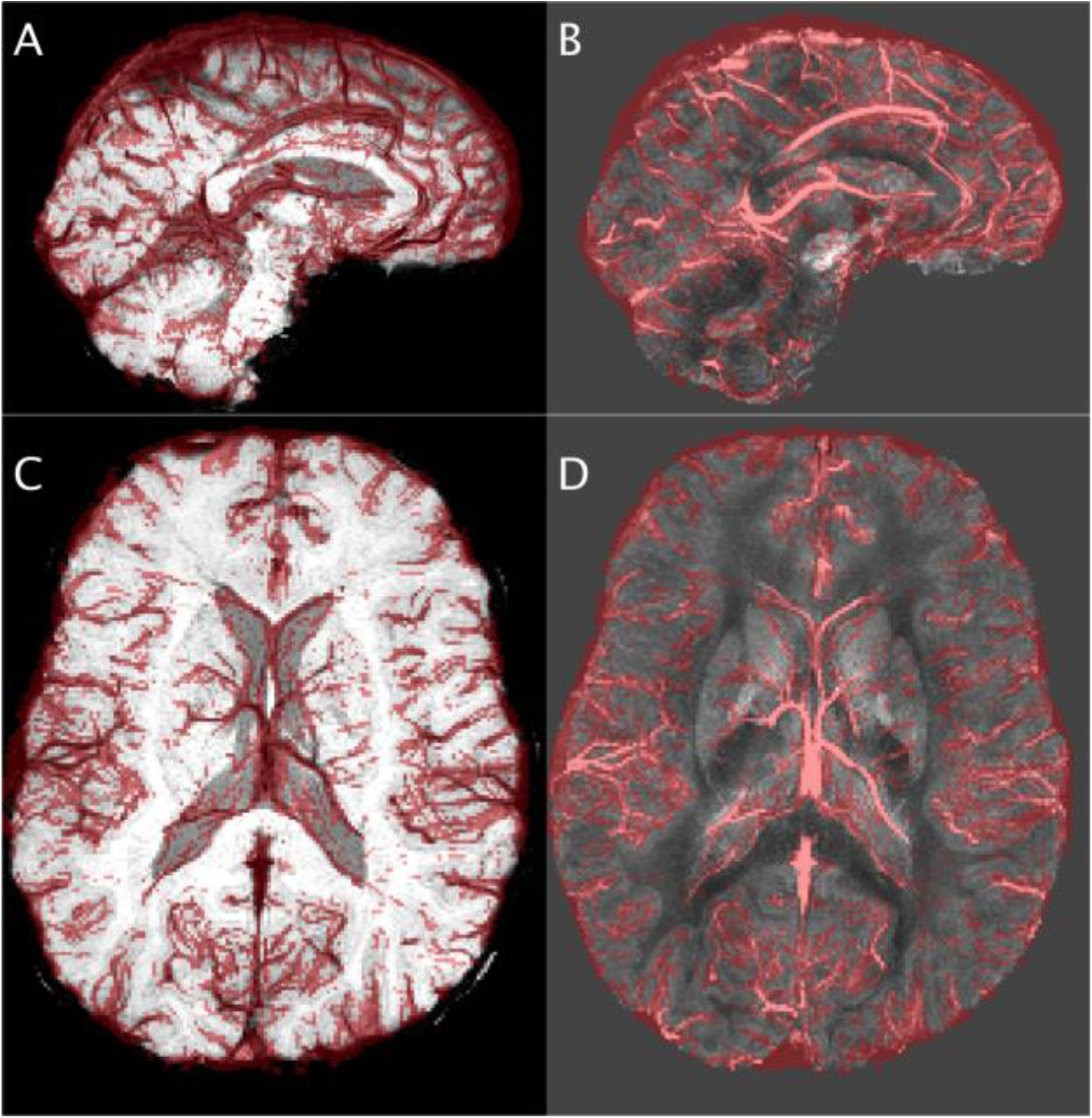
Example images of the manual vein tracings. Minimum-intensity projections for SWI (A and C) and maximum-intensity projections for QSM (B and D), with a semi-transparent vein mask overlay in red. Sagittal (9mm slab, A and B) and axial (18mm slab, C and D) slices are shown from a single subject.

### 2.3 Composite vein image process

The composite vein (CV) image (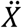) was constructed by combining a vein atlas, with a SWI image and a QSM image. The three inputs are referred to as *C*, where *C* = {*atlas* = 1, *SWI* = 2, *QSM* = 3}, and are denoted *X_c_*.

The CV method combines the three inputs with weights based on the relative predictive power of each input in different regions of the brain. The relative predictive power was captured in a *template prior* (*P_c_*) for each of the three inputs (Figure **2A**). The inputs were normalized 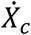, and then combined using a weighted-mean with the weights derived from the priors, *P_c_*.
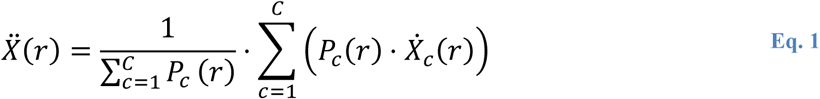

**Figure 2.**
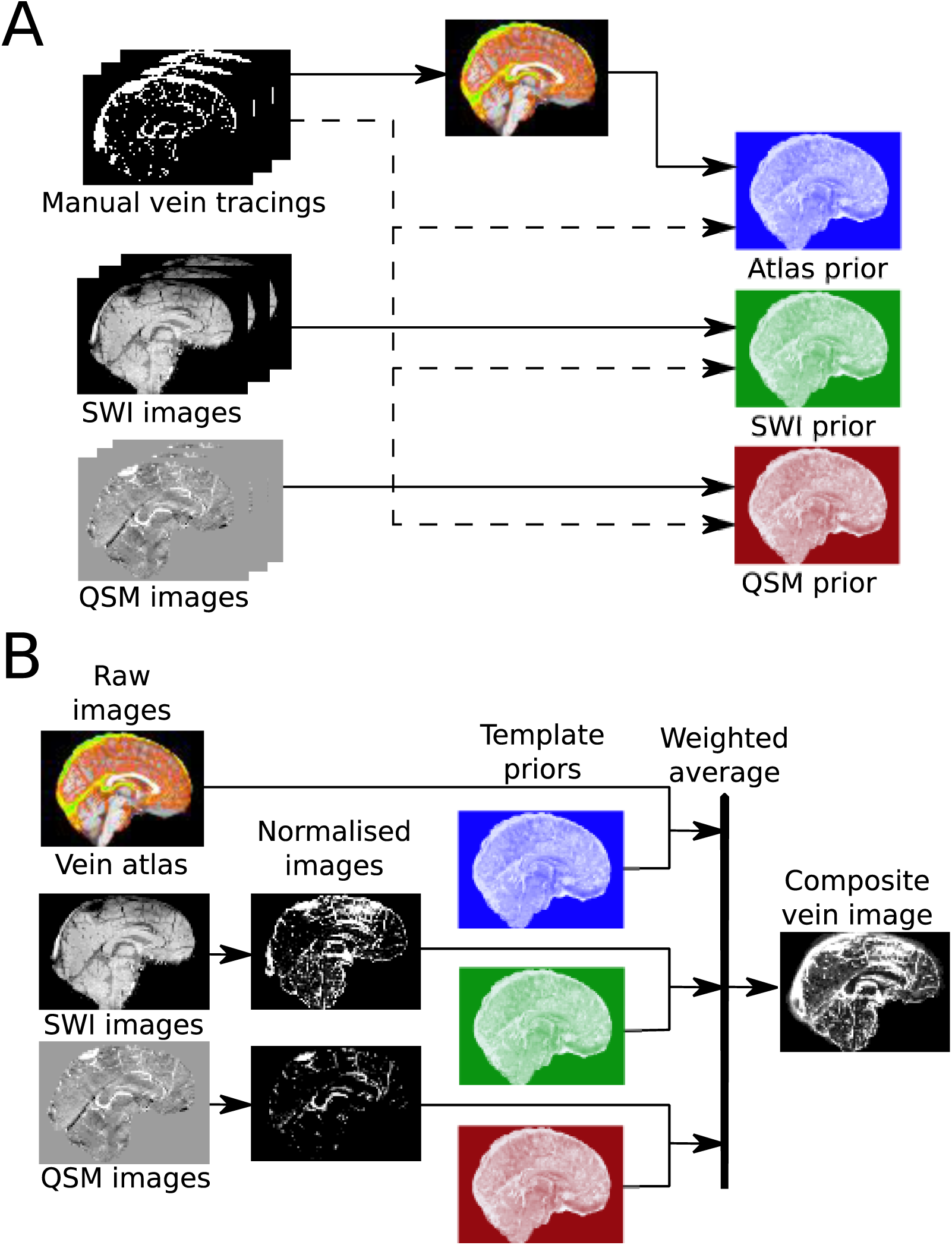
Schematic describing data flow and process for training the priors and atlas (A) and producing a composite vein image (B). The training data sets (manual vein tracings, SWI images, QSM images) in (A) are used to calculate the vein atlas and template priors. Operations in (A) occur within MNI template space. The vein atlas, atlas prior, SWI prior and QSM prior in (A) are inputs to (B) once they are interpolated into subject space. The training data set in (A) does not include the subject images in (B) to ensure independence.

The CV image, 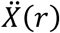, for each subject used the atlas and priors pre-calculated from the training cohort (*M*) (Figure **2A**). For each subject in this study, the training cohort consisted of the other nine subjects to ensure independence of the atlas and priors for each subject specific CV image.

The atlas construction, normalization process, and details of the template priors are explained in the following sub-sections.

### 2.4 Vein atlas

Manually traced vein masks (*Y*) were weighted to reflect the variance and uncertainty of the human observer (*W*).
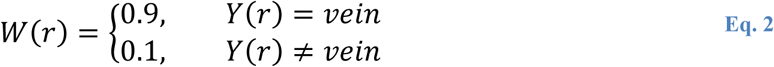

The weighted tracings (*W_i_*) for each subject, *i*, were interpolated into 0.5mm MNI space (Montreal Neurological Institute standard brain atlas) and the average (mean) calculated for each voxel, *r*, to construct a vein atlas.
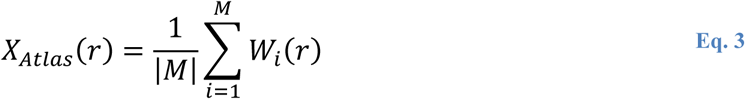

Visual inspection of the atlas was performed to explore the variability of vein location between subjects within the cohort.

### 2.5 Normalisation

The SWI and QSM images were processed separately to remove biases and to normalize their voxel intensities using a Gaussian mixture model (GMM) with two components (vein and non-vein). This approach has been explored previously (Ward et al., 2015, 2017b), and similar techniques have been used in blood vessel segmentation before (Bazin et al., 2016; Bériault et al., 2015, 2014; Hassouna et al., 2006).

For each image (SWI and QSM), a GMM was fit using a log-likelihood expectation-maximisation approach (Dempster et al., 1977). Both GMMs (one for SWI and one for QSM) used the same initial seed taken from the QSM images (*X_QSM_* > 0.05*ppm*) to impart prior knowledge of the components. High-pass filtering was also applied to *X_SWI_*.

The GMM process mapped the image contrasts to a unity space [0,1] and reduced the presence of subject specific biases. Normalized images (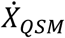 and 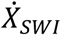) were produced from the mixture coefficient of the vein component for each voxel.
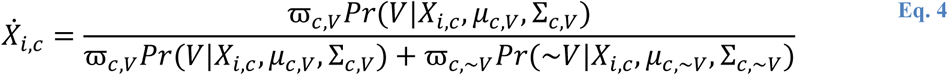

where *c* was the image (SWI or QSM), *i* was the voxel, and *Pr*(*V* | *X_c_*, *μ_c_*_,*V*_, Σ_c,V_) was the posterior probability of being labelled vein (or not vein, ~*V*) given the distribution parameters for the image-specific GMM: *μ*, Σ, relative abundance ϖ, and the voxel intensity, *X*. Supporting information on this process has been published previously (Ward, 2017).

The atlas (*X_Atlas_*) is intrinsically normalized, and was interpolated into the subject space to provide the final input (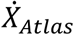).

### 2.6 Template priors

Subject specific confidence maps, *p_i,c_*, were calculated using log-loss scoring (Dowe, 2008; Good, 1952) for each input (*c* ∈ *C*) using Eq. 5. Voxel location (*r*) and subject (*i*) have been omitted from this equation for clarity.
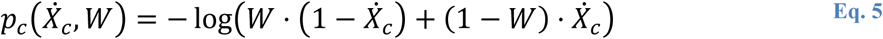

The template priors (*P_c_*) for SWI, QSM and the vein atlas were calculated by taking the cohort mean of the confidence maps interpolated into MNI space.
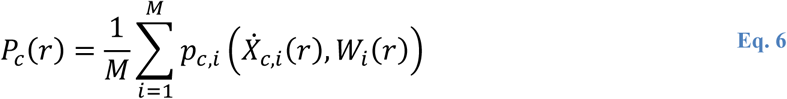

The predictive power represented in the priors for each information source (*P_Atlas_*, *P_SWI_*, and *P_QSM_*) was examined visually. In order to visualize all three values, each was normalized, 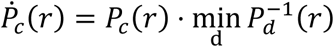, and encoded in a colour channel of a colour image (blue, green, red for *P_Atlas_*, *P_SWI_*, and *P_QSM_* respectively).

### 2.7 Performance evaluation

Vein contrast in the CV image was assessed in comparison to SWI and QSM images using automated vein segmentation techniques. Three vein segmentation techniques were employed to compute venograms for each of the three image sets (the CV images, SWI images and QSM images). The first segmentation technique was a Hessian-based vesselness filter followed by an Otsu threshold (VN) (Frangi et al., 1998; Otsu, 1975). The second was a statistical method based on an Ising model Markov random field using an anisotropic graph (MRF) (Bériault et al., 2014). The third was a recursive ridge-based filter (RR) (Bazin et al., 2016).

The accuracy of the venograms from each image set was evaluated with standard metrics (Table **1**). Many standard overlap metrics are not informative due to the high surface-to-area ratio of cerebral vein masks. To overcome this limitation, dilated versions of many of the metrics were used (Bazin et al., 2016). The metric values for the SWI and QSM image based venograms were calculated as benchmarks, and the differences between these reference values and the venograms computed from the CV image were examined.

**Table 1.** Venogram accuracy metrics. The metrics compare the manual mask of veins (***V***) and non-veins (***N***) with an automated estimate (***V***’ and ***N***’). A mask morphologically dilated by one-voxel is preceded by a ***δ***, e.g. ***δV*** is the dilated manual vein mask. Dilation was used as the boundaries of small vein may be uncertain due to bloom and partial volume. The Hausdorff distance (41), 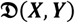, is the mean value in a minimum distance map, i.e., the minimum distance from each surface voxel in mask ***X*** to a surface voxel in mask ***Y***. The surface is defined as all voxels removed by a one-voxel morphological erosion.

| Name | Equation |
| --- | --- |
| Number of true-positives (veins) | $TP = \vert V \cap V'\vert $ $\delta TP = \frac{1}{2} (\vert \delta V \cap V'\vert + \vert V \cap \delta V'\vert )$ |
| Number of true-negatives (non-veins) | $TN = \vert N \cap N'\vert $ $\delta TN = \frac{1}{2} (\vert \delta N \cap N'\vert + \vert N \cap \delta N'\vert )$ |
| Number of false-positives | $FP = \vert N \cap V'\vert $ |
| Number of false-negatives | $FN = \vert V \cap N'\vert $ |
| Accuracy | $ACC = \frac{\delta TP + \delta TN}{\vert V \cup N\vert }$ |
| Sensitivity | $SE = \frac{\vert \delta V' \cap V\vert }{\vert V\vert }$ |
| Specificity | $SP = \frac{\vert \delta N' \cap N\vert }{\vert N\vert }$ |
| Positive-predictor value | $PPV = \frac{\vert \delta V \cap V'\vert }{\vert V'\vert }$ |
| Negative-predictor value | $NPV = \frac{\vert \delta N \cap N'\vert }{\vert N'\vert }$ |
| Dice similarity score (Dice, 1945) | $DSS = \frac{2 \cdot \delta TP}{\vert V\vert + \vert V'\vert }$ |
| Matthews correlation coefficient (Matthews, 1975) | $MCC = \frac{TP \cdot TN - FP \cdot FN}{\sqrt{(TP + FP)(TP + FN)(TN + FP)(TN + FN)}}$ |
| Mean Hausdorff distance (Shonkwiler, 1989) | $MHD = \frac{1}{2} (\mathfrak{D}(V, V') + \mathfrak{D}(V', V))$ |
| Average volume difference | $AVD = \frac{\vert FP - FN\vert }{\vert V\vert }$ |

Two techniques had parameters that required training (VN and MRF). A leave-one-out approach to training was taken, with the performance of the left-out subject recorded for comparison purposes. The parameters (*Θ*) were optimized in a standardized space to minimize a composite cost function (Z), which included a regularization term (*Λ*). 
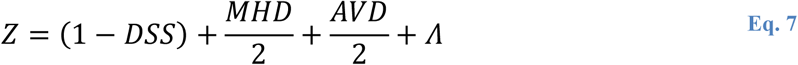

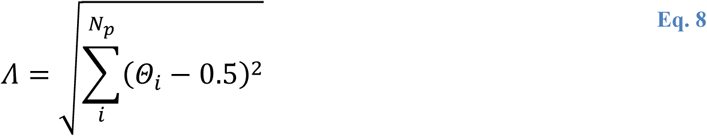

where *N_p_* is the number of parameters to train. Dice similarity score (DSS) (Dice, 1945), mean Hausdorff distance (MHD) (Shonkwiler, 1989) and average volume difference (AVD) are described in Table **1**. Scaling factors were chosen to normalize metric ranges as DSS is constrained to a unity range, whilst MHD and AVD are unbounded.

A gradient descent algorithm was used to search the parameter space from an initial parameter estimate. The initial parameter estimate was the optimal parameter values from a uniform random sample of the entire parameter space (1000 samples). Each iteration of the search algorithm used 32 randomly sampled potential steps. Samples were taken from a hypercube with dimensions equal to 5% of the parameter space. In all cases the cost function was found to have converged before 50 iterations. All parameters are reported in Table **2**. All operations were performed in MATLAB 2015b using the MASSIVE supercomputer (Goscinski et al., 2014).

**Table 2.** Tuned parameter descriptions for each automated segmentation technique. The trained value represents the mean and standard deviation across the 10 parameter sets trained.

| Technique | Parameter | Description | Trained value |  |  |
| --- | --- | --- | --- | --- | --- |
|  |  |  | CV | SWI-Only | QSM-Only |
| MRF | $\beta_{ratio}$ | Weighting factor between vein and non-vein in neighbours in clique potential calculation. | $0.10 \pm 0.02$ | $0.58 \pm 0.02$ | $0.48 \pm 0.01$ |
| VN | $\alpha$ | Non-plane like factor | $0.24 \pm 0.01$ | $0.19 \pm 0.02$ | $0.61 \pm 0.13$ |
| | $\beta$ | Non-blob like factor | $0.77 \pm 0.09$ | $0.34 \pm 0.21$ | $0.16 \pm 0.02$ |
| | $\gamma$ | Intensity factor | $1.61 \pm 0.08$ | $3.89 \pm 0.22$ | $7.65 \pm 1.23$ |
| | scale | Maximum scale explored | $1.00 \pm 0.01$ | $1.00 \pm 0.01$ | $1.00 \pm 0.01$ |
| RR | Recommended parameters specified by creator suitable for contrasts used (none trained). |  |  |  |  |

Ten sets of parameters were trained for each technique (Table **2**), using a different subset of nine subjects from the ten available. The mean and standard deviation of the results are shown in Table **2**. The difference in cost function value between the mean of the training set (nine subjects) and the left-out validation subject was low. In the majority of cases the validation score was better than the worst individual score in the training set.

The difference in performance was assessed between the CV images and SWI images, and CV images and QSM images. The magnitude of performance difference was quantified using Cohen’s d (Cohen, 2013) and interpreted on a qualitative scale (Sawilowsky, 2009). The significance of the difference was tested using a paired two-tailed Wilcoxon signed-rank test (Wilcoxon, 1945). A description of all metrics used can be found in Table **1**.

## 3. Results

All permutations of vein segmentation technique, performance metric and benchmark image (SWI or QSM) were explored, resulting in 60 comparisons. When using the CV image, 77% of the permutations showed a large or higher improvement (Cohen’s d > 0.80) that was statistically significant (p<0.05), compared to a negative effect in 5% of the permutations. The mean effect size across all permutations was 1.1 (large) and very large effects in favor of the CV image were found in 65% of comparisons. The results are shown pictographically in Figure **3**.

**Figure 3.**
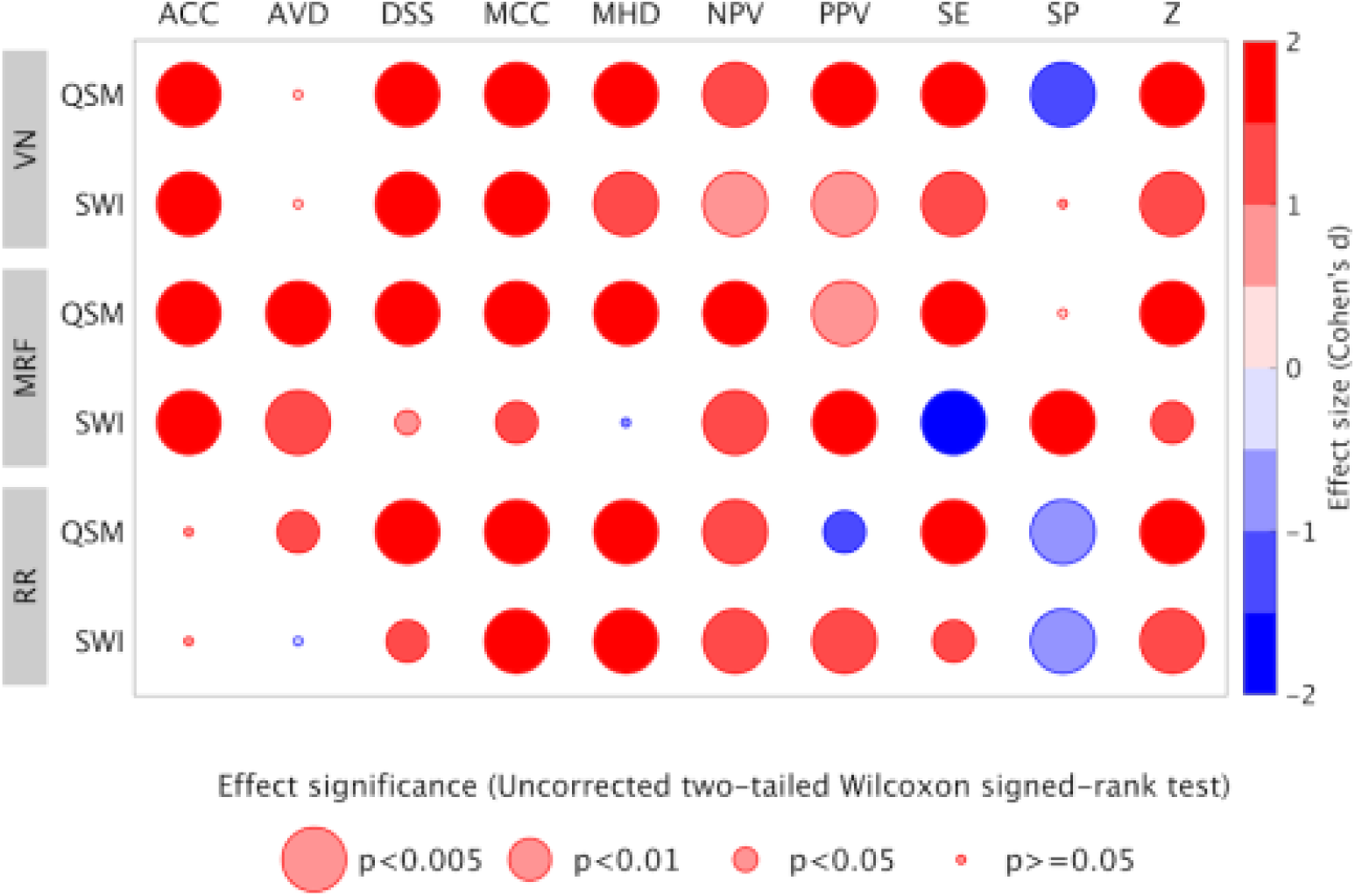
The improvement in performance of three automated segmentation techniques when using the CV image compared to two alternative images. The size of the improvement (Cohen’s d) and the statistical significance of the improvement (p-values) are displayed in colour and size respectively. Red circles indicate superior performance using the CV image. Large circles indicate more significant results. Each row corresponds to an automated segmentation technique (RR, MRF, or VN) and input image (SWI or QSM). Each column denotes a different performance metric. All metric abbreviations are provided in Table 1. Statistical significance was measured using a two-tailed Wilcoxon signed-rank test and is uncorrected.

Negative results (5%) occurred in paired-metrics and were not observed in comprehensive balanced metrics, such as DSS, MCC and MHD, which overwhelmingly displayed performance improvements with the CV image. In paired-metrics, a corresponding positive effect was found for each negative effect in the metric pair, such as negative specificity and positive sensitivity for the QSM image when using the VN technique. In these paired cases the effect size was comparable for both positive and negative results.

Inconclusive results were most common for average-volume difference (AVD) (50% of comparisons). AVD measures bias in the errors of the final masks, rather than a direct measure of performance. Greater improvement was shown in comparison to the QSM based venograms relative to SWI based venograms.

The similarity of vein locations between subjects was represented in the vein atlas (Figure **4**). High atlas values indicated consistent vein locations between subjects. The highest value voxels were found near the major veins, including in the superior sagittal sinus, dural sinuses, straight sinus and internal cerebral veins (green regions in Figure **4**). The deep gray matter structures, the inferior frontal and inferior temporal regions showed lower values.

**Figure 4.**
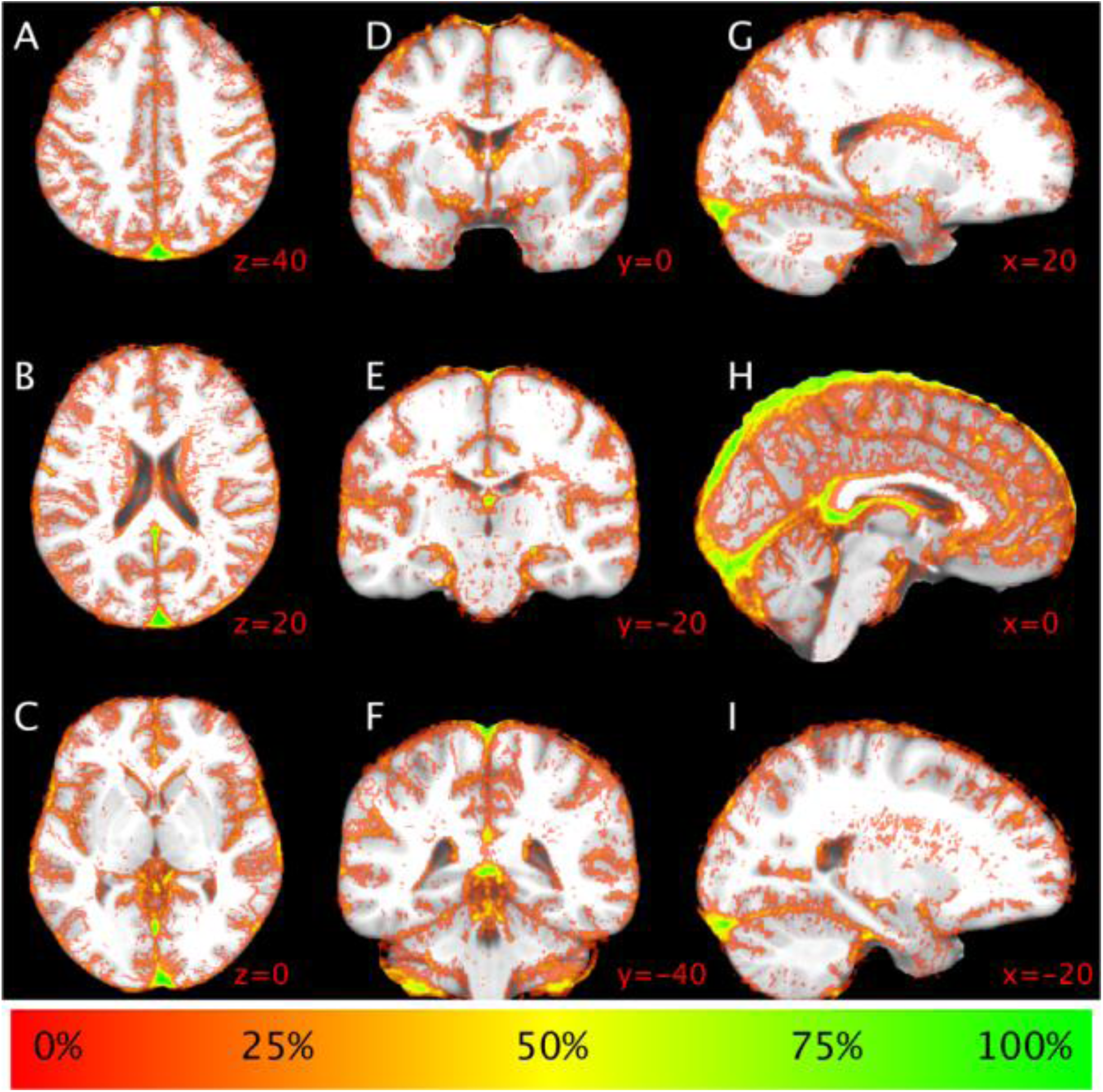
Vein frequency atlas (***X_Atlas_***) demonstrating high reproducibility in vein location across subjects in the major veins (e.g. sagittal sinus, green/yellow) and low reproducibility in the deep grey matter structures. Values of 0% are transparent. Slice coordinates (red) are in NMI atlas space, with axial projections (A-C), coronal projections (D-F) and sagittal projections (G-I).

The relative predictive power of the atlas, SWI and QSM template priors was observed to be heterogeneous across brain regions (Figure **5**). QSM was found to have comparatively higher power in the falx cerebri and lower power in the deep-gray matter structures, relative to SWI and the atlas. The atlas was highest in the deep-gray matter, particularly on the edge of structures. SWI had higher predictive power on the superior surface of the cortex, and lower power on the inferior surface of the brain.

**Figure 5.**
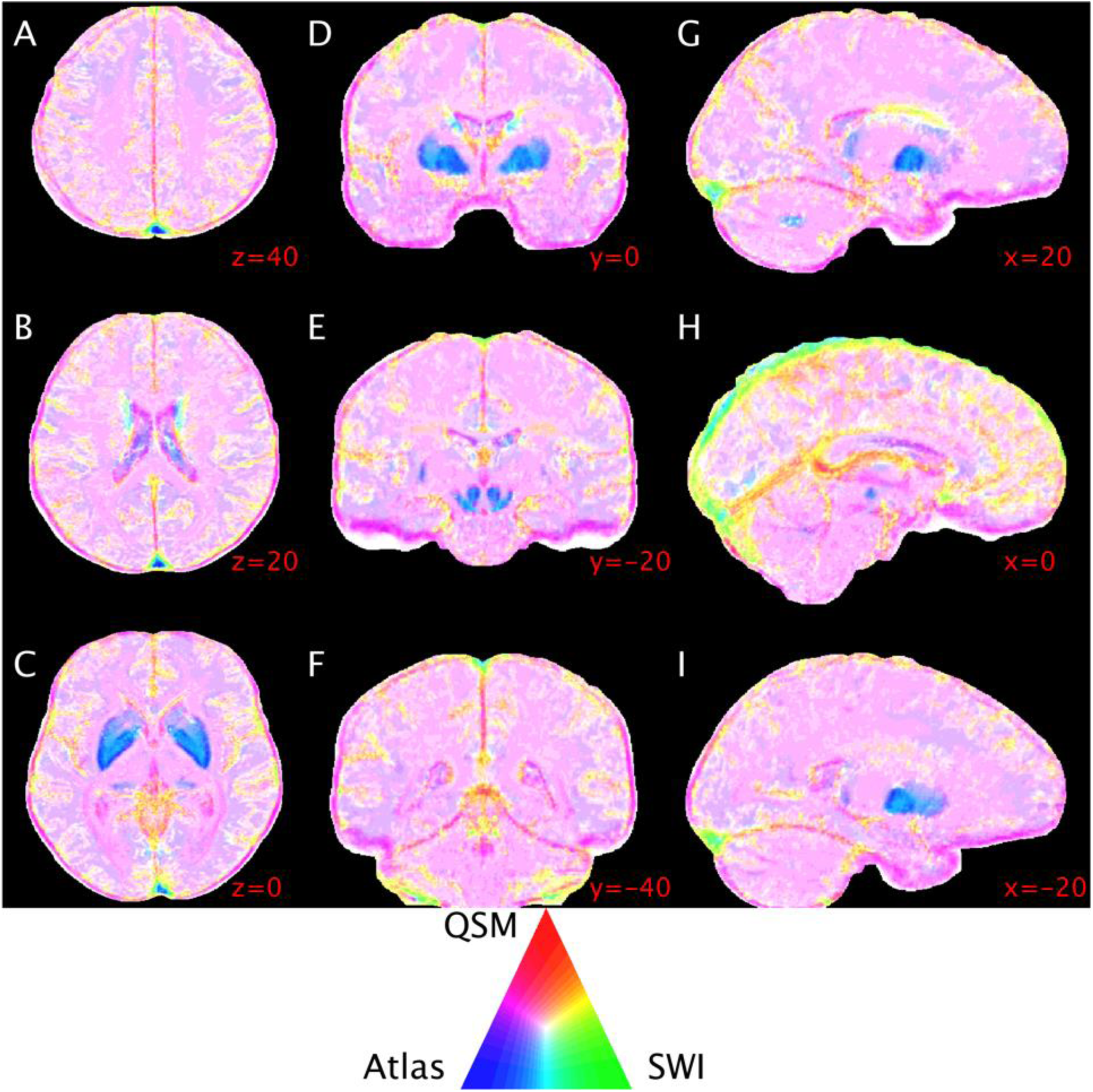
Slices from the atlas, SWI and QSM template priors (***P_Atlas_***, ***P_SWI_***, and ***P_QSM_*** respectively), color-coded to represent the relative weights of each. Blue regions show where ***P_Atlas_*** is highest, green regions where ***P_SWI_*** is highest and red regions where ***P_QSM_*** is highest. A triangular colour-map is included. Slice coordinates (red) are in MNI atlas space, with axial projections (A-C), coronal projections (D-F) and sagittal projections (G-I).

## 4. Discussion

In this work a composite vein (CV) imaging technique was proposed that combined three sources of vein information, an atlas, a SWI image, and a QSM image. The CV image showed a large improvement in vein segmentation accuracy when compared with SWI and QSM images. A robust improvement was observed in the majority of permutations across ten performance metrics and three segmentation methods.

The CV image was found to combine the complementary strengths of SWI and QSM, and produce an image with significantly improved vein contrast by incorporating the relative predictive power of SWI and QSM in a weighted-average approach. A comprehensive analysis was performed using over 1000 MRI slices, including anisotropic and isotropic acquisitions, multiple automated segmentation techniques, and multiple performance metrics.

The template priors characterized the anatomically heterogeneous value of the three inputs. The atlas template prior (*P_Atlas_*) had higher relative values to SWI and QSM in the deep gray matter structures possibly due to non-venous iron deposits in tissue. However, the relative predictive power of SWI increased towards the center of these structures. High-pass filtering in SWI may be the cause of this effect by reducing the low-frequency spatial contrast of these iron sources and increasing the sharpness of structure boundaries. A common trend observed in the larger veins, particularly those in the interhemispheric region, was higher SWI predictive power in the center and higher QSM predictive power at the vessel wall, extending into the surrounding tissue. Extravascular enhancement on SWI images may be the source of the decrease in SWI predictive power at the vessel wall. Two exceptions to the greater predictive power of SWI in the center of larger veins were the superior sagittal sinus and the transverse sinus, where the predictive power of the atlas was higher. Decreased anatomical variability and hyper-intense GRE signal due to imperfect flow compensation are possible causes of the reduction in SWI predictive power relative to the atlas.

There was minimal discrepancy between performance of the training and validation datasets. The tuned parameter values for each of the ten-independent datasets had low variance (Table **2**) despite the stochastic nature of the initial parameter values. These two results indicate minimal over-fitting occurred when optimizing the technique parameters.

Comparative analysis of published automated vein segmentation techniques is difficult for a number of reasons. Manually traced ground truth vein masks typically cover small manually traced regions and/or minimum-intensity projections (Bazin et al., 2016; Bériault et al., 2014; Monti et al., 2015). The small regions may not be indicative of performance across the entire brain, and are a source of variability between studies that cannot be controlled for. Studies also use different performance metrics including sensitivity and specificity (Monti et al., 2015), positive-predictor value, negative predictor value and overlap (Bazin et al., 2016), and accuracy (Bériault et al., 2014). The use of different metrics may be due to the specific application that each technique has been designed for. However, the selective use of metrics can result in one-sided conclusions being drawn, and can frustrate meta-analysis efforts.

The quantification of vein segmentation performance is a contentious issue (Gerig et al., 2001). To capture compensatory behavior, where one metric is optimized at the expense of another, metrics are often reported in quasi-orthogonal pairs, such as specificity and sensitivity. Although, when one label (vein) is less numerous in abundance than the other (non-vein) the trade-off between pairs will not be even due to the disparate magnitude of the denominators. Non-paired metrics, such as overlap and volume difference metrics, are more robust in these scenarios, albeit at the expense of interpretability. Comprehensive reporting of multiple metrics should be adopted to enhance both interpretability and transparency.

The template priors, vein atlas, and manual vein tracings are publicly available (Ward et al., 2016a). The manual vein tracing required hundreds of hours to complete, and the release of the segmented data may facilitate future work in vein segmentation techniques. The data sharing may result in a large, publicly available set of cross-validated ground truth vein images for future collaborative studies.

### 4.1 Limitations

A distinction should be considered between accurate vein segmentation and accurate imitation of manually traced vein masks. The tracings are not a direct measure of veins, but a subjective radiological interpretation (Drew et al., 2013), and are produced in the presence of the artifacts that occur in susceptibility-based MRI. A ground truth that is independent of MRI-based artifacts would be required to directly quantify vein segmentation accuracy.

Furthermore, only binary masks have been addressed in this work. However, vein geometry does not conform to a cubic grid. Recent work has found significant error associated with binary representations of veins in simulated QSM images (Ward et al., 2017a). Further work will explore non-binary manual tracings to incorporate both partial volume and marker uncertainty.

## 5. Conclusions

A large improvement in venogram accuracy was achieved using the composite vein image technique. The composite vein image technique incorporates the heterogeneous vein contrast profile across the brain to extract the complementary information available from SWI and QSM images, and a vein atlas. The technique’s performance was evaluated with multiple segmentation techniques and metrics. The accuracy provided by the composite vein image allows improved quantification of cerebrovenous topology and cerebrovascular oxygenation using MRI.

## Acknowledgements

We thank the investigators of the ASPREE study and the ASPREE-NEURO sub-study for the coordination and recruitment of the volunteers. We thank the volunteers for their time and willingness to participate.

The Alzheimer’s Australia Dementia Research Foundation (AADRF), the Victorian Life Sciences Computation Initiative (VLSCI), the Multi-model Australian ScienceS Imaging and Visualisation Environment (MASSIVE) and the National Health and Medical Research Council (NHMRC) supported this work (NHMRC Grant APP1086188).

We would like to thank Dr. Pierre-Louis Bazin and Dr. Silvain Bériault for their assistance in implementing and applying their vein segmentation methods.

Initial work and preliminary findings related to this study were previously presented at annual meetings of the International Society for Magnetic Resonance in Medicine (Ward et al., 2015, 2016b, 2017d).

